# Penetrance and pleiotropy of polygenic risk scores for schizophrenia in 106,160 patients across four healthcare systems

**DOI:** 10.1101/421164

**Authors:** Amanda B Zheutlin, Jessica Dennis, Richard Karlsson Linnér, Arden Moscati, Nicole Restrepo, Peter Straub, Douglas Ruderfer, Victor M Castro, Chia-Yen Chen, Tian Ge, Laura M Huckins, Alexander Charney, H Lester Kirchner, Eli A Stahl, Christopher F Chabris, Lea K Davis, Jordan W Smoller

## Abstract

**OBJECTIVE:** Individuals at high risk for schizophrenia may benefit from early intervention but few validated risk predictors are available. Genetic profiling is one approach to risk stratification that has been extensively validated in research cohorts, but its utility in clinical settings remains largely unexplored. Moreover, the broad health consequences of a high genetic risk of schizophrenia are poorly understood, despite being relevant to treatment decisions.

**METHOD:** We used electronic health records for 106,160 patients from four healthcare systems to evaluate the penetrance and pleiotropy of genetic risk for schizophrenia. Polygenic risk scores (PRSs) for schizophrenia were calculated from summary statistics and tested for association with 1359 disease categories, including schizophrenia and psychosis, in phenome-wide association studies. Effects were combined through meta-analysis across sites.

**RESULTS:** PRSs were robustly associated with schizophrenia (odds ratio per standard deviation increase in PRS=1.55 [95% confidence interval (CI), 1.4-1.7], *p*=4.48 x 10^-16^) and patients in the highest risk decile of the PRS distribution had up to 4.6-fold increased odds of schizophrenia compared to those in the bottom decile (95% CI, 2.9-7.3, *p*=1.37 x 10^-10^). PRSs were also positively associated with a range of other phenotypes, including anxiety, mood, substance use, neurological, and personality disorders, as well as suicidal behavior, memory loss, and urinary syndromes; they were inversely related to obesity.

**CONCLUSIONS:** We demonstrate that an available measure of genetic risk for schizophrenia is robustly associated with schizophrenia in healthcare settings and has pleiotropic effects on related psychiatric disorders as well as other medical syndromes. Our results provide an initial indication of the opportunities and limitations that may arise with the future application of PRS testing in healthcare systems.

## Introduction

Psychiatric disorders are common and responsible for an enormous burden of suffering^1,2^. Approximately 18% of individuals globally suffer from mental illness every year^3^, 44.7 million of whom live in the US^4^. Early detection and intervention for serious mental illness is associated with improved outcomes^5–8^. However, few reliable predictors of risk or clinical outcomes have been identified. Given the substantial heritability of many psychiatric disorders^9^ and their polygenic architecture^10^, there is increasing interest in using quantitative measures of genetic risk for risk stratification^11^. Polygenic risk scores (PRSs), in particular, are easy and inexpensive to generate and can be applied well before illness onset, making them a promising candidate for clinical integration^12^. In fact, a recent study investigating the clinical utility of PRSs for several common, non-psychiatric diseases found that PRS can identify a larger fraction of high risk individuals than are identified by clinically-validated monogenic mutations, and called explicitly for evaluations of these scores in clinical settings^13^.

To date, PRSs for neuropsychiatric disorders have primarily been evaluated in highly ascertained research samples. Typically, cases have obtained a diagnosis through lengthy clinician interviews and controls have no psychiatric history (“clean” cases and controls). In order to bring PRSs to the clinic, however, they must first demonstrate association with diagnoses in real-world clinical settings, where data are often much messier. Among psychiatric disorders, schizophrenia is perhaps the best candidate for future clinical integration of PRS profiling as it is highly heritable, has the best performing PRS among psychiatric disorders in terms of proportion of phenotypic variance explained (7%)^14^, and can be difficult to distinguish from other psychiatric conditions that involve psychosis and mood disturbance. Accordingly, we selected the schizophrenia PRS for the present study as it is the most viable test case for eventual clinical validation of a psychiatric PRS.

We recently established the PsycheMERGE consortium within the NIH-funded Electronic Medical Records and Genomics (eMERGE) Network^15, 16^ to leverage electronic health record (EHR) data linked to genomic data to facilitate psychiatric genetic research^17^. In this first report from PsycheMERGE, we evaluated the performance of a schizophrenia PRS generated from summary statistics published by the Psychiatric Genomics Consortium^14^ using EHR data on more than 100,000 patients from four large healthcare systems (Geisinger Health System, Mount Sinai Health System, Partners Healthcare System, and Vanderbilt University Medical Center). We assessed the relative and absolute risk for schizophrenia among individuals at the highest level of genetic risk and considered the clinical utility of the PRS for risk stratification. We also examined pleiotropic effects of the schizophrenia PRS with real-world clinical data by conducting a phenome-wide association study (PheWAS) of 1359 disease categories. To our knowledge this is the first effort to combine PheWAS effects across multiple hospital-based biobanks.

Finally, we conducted follow-up analyses to characterize the nature of the pleiotropic effects of the schizophrenia PRS. Cross-phenotype associations of polygenic liability to schizophrenia may occur in at least two scenarios^18^. In the first (“biological pleiotropy”), the PRS contributes independently to multiple phenotypes. In the second scenario (“mediated pleiotropy”), the PRS increases liability to a second disorder that occurs as a consequence of schizophrenia itself. For example, an association between schizophrenia polygenic risk and diabetes could occur because individuals diagnosed with schizophrenia are more likely to have both elevated schizophrenia PRS *and* to be prescribed antipsychotic medications which may result in weight gain and increased liability to diabetes. In this case, the observed relationship between schizophrenia risk and diabetes is mediated by the use of antipsychotic medication. These scenarios may be difficult to completely disentangle. However, here we use individual-level EHR data to determine whether associations with genetic risk for schizophrenia persist after conditioning on a clinical diagnosis of schizophrenia, related psychosis, or prescription of antipsychotic medications.

## Methods

### Hospital-based Biobanks

Patients that consented to participate in one of four large healthcare system-based biobanks – the MyCode Community Health Initiative at the Geisinger Health System (GHS)^19^, the BioMe Biobank at the Mount Sinai School of Medicine (MSSM)^20^, the Partners Healthcare System (PHS) biobank^21^, or the Vanderbilt University Medical Center (VUMC) biobank (BioVU)^22^ – and had available EHR and genotype data were included in these analyses. At each site, patients were recruited from the general healthcare system population without systematic recruitment for any particular disease or diagnosis. It is well known that PRS calculated from GWAS performed primarily in one ancestry demonstrates poorer performance in other ancestries as a function of differing LD structures with causal variants and lack of diversity on genotyping platforms^23, 24^. Thus, this study was limited to patients of European-American ancestry with genetic data that met standard quality control thresholds (see Quality Control of Genetic Data). Besides these data availability and ancestry filters, no further inclusion or exclusion criteria were applied. Our final sample included 44,436 patients from GHS, 9,569 patients from MSSM, 18,461 patients from PHS, and 33,694 patients from VUMC (106,160 total participants). All patients gave informed consent for biobank research for which IRB approval was obtained at each site.

### Quality Control of Genetic Data

Samples were genotyped, imputed, and cleaned at each site individually, the details of which are described in Supplementary Methods. However, quality control procedures at each site followed a similar standard pipeline. DNA from blood samples obtained from biobank participants were assayed using Illumina bead arrays (OmniExpress Exome, Global Screening, MEGA, MEGA^EX^, or MEG BeadChips) containing approximately 700,000 to two million markers. Samples at each site were genotyped in multiple batches; indicators for genotyping platform and batch were included as covariates in the analyses. As described in Supplementary Methods, single nucleotide polymorphisms (SNPs) were excluded using filters for call rate, minor allele frequency, and heterozygosity at a minimum. Individuals were excluded for excessive missing data or sex errors; a random individual from any pair of related individuals was also excluded (pihat > .2). Principal components or self-reported ancestry was used to identify individuals of European ancestry. SNPs that passed the initial phase of quality control were imputed and then converted to best-guess genotypes where only high-quality markers were retained. Ten principal components were generated within the European sample to use as ancestry covariates in all subsequent analyses.

### Polygenic Risk Scores

In order to quantify genetic risk for schizophrenia, we calculated PRSs using summary statistics from the Psychiatric Genomics Consortium genome-wide association study (GWAS) of schizophrenia^14^, which included odds ratios (ORs) for 9,444,230 variants. PRSs were calculated via two methods, a simple and widely used approach where SNPs are pruned based on linkage disequilibrium (LD) and association p-values, and a Bayesian approach that can increase accuracy by directly modeling LD structure and adjusting SNP weights accordingly.

#### LD-pruned PRS

We excluded rare variants (minor allele frequency <1%) and variants on the X chromosome and then, at each site, clumped SNPs based on association p-value (the variant with the smallest p-value within a 250kb range was retained and all those in LD, *r^2^* > .1, were removed). The resulting SNP lists included 146,464 at GHS, 79,837 at MSSM, 166,477 at PHS, and 229,355 variants at VUMC. Using all available variants (i.e., using a p-value threshold of 1.0 for inclusion), we generated PRSs for each individual by summing all risk-associated variants weighted by the log(OR) for that allele from the GWAS. PRSs were converted to z-scores within each healthcare system to standardize effects across all sites. LD pruning and PRS generation were done using PRSice^25^.

#### Bayesian PRS

We used PRS-CS, a Bayesian polygenic prediction method, as an alternative approach for PRS calculation. PRS-CS places a continuous shrinkage (CS) prior on SNP effect sizes and infers posterior SNP weights using GWAS summary statistics and an external LD reference panel (1000 Genomes Project European samples; N=503). PRS-CS enables multivariate modeling of local LD patterns and is robust to diverse underlying genetic architectures, and thus can increase the accuracy of PRS over conventional approaches^26^. At each site, weights for all imputed SNPs present on the 1000 Genomes reference panel and HapMap3 panel were estimated using PRS-CS, resulting in 833,502 available SNPs at GHS, 971,463 at MSSM, 833,502 at PHS, and 604,645 at VUMC. The global shrinkage parameter in the CS prior was fixed at 1 to reflect the highly polygenic genetic architecture of schizophrenia. We generated PRSs for each individual by summing all risk-associated variants weighted by the posterior effect size inferred by PRS-CS for that allele and then converted PRSs to z-scores within each healthcare system. A Python package for PRS-CS is available on GitHub repository (https://github.com/getian107/PRScs). PRSs were calculated using PLINK 1.9^27^.

### EHR-derived Phenotypes

EHRs contain thousands of diagnostic billing codes from the International Classification of Diseases, 9^th^ and 10^th^ editions (ICD-9/10) which are arranged hierarchically. For example, ICD9:295 is ‘schizophrenic disorders’, ICD9:295.1 is ‘disorganized type schizophrenia’, and ICD9:295.12 is ‘disorganized type schizophrenia, chronic state’; in total, the ICD9:295 category contains 71 individual ICD-9 codes. To define case status for a variety of diseases, we extracted all ICD-9 and ICD-10 codes available for participating subjects and grouped codes into 1860 disease categories (called ‘phecodes’) using a hierarchical structure previously developed and validated^28, 29^. For “schizophrenia and other psychotic disorders”, for example, 89 individual ICD-9 codes – all 71 ICD9:295 codes and 18 related codes (e.g., 298.9, unspecified psychosis) – and 22 ICD-10 codes were mapped to this disease category.

Cases and controls were designated for each phecode. Individuals with two or more relevant ICD-9/10 codes were considered a case, those with zero relevant codes were considered a control, and individuals with only one code were excluded^30^. To enable analyses of phenome-wide diagnoses that may have varying ages of onset, we did not restrict the age range of participants. The proportion of patients (cases and controls) included in a given PRS-phecode association varied depending on the prevalence of single-code individuals, but the median was 98%-100% at each site. Phecodes for which there were fewer than 100 cases were excluded from the PheWAS.

### Statistical Analyses

#### Penetrance of schizophrenia PRS in healthcare systems

To assess the penetrance of schizophrenia PRS, we measured absolute risk (case prevalence as a function of PRS) and relative risk (ORs for the top decile of schizophrenia PRS relative to the remaining population, as well as the bottom decile) for schizophrenia and psychotic disorders. ORs were calculated at each site for both PRS methods (LD-pruned and Bayesian), regardless of the number of available cases, and then the log(OR)s were combined through fixed-effect inverse variance-weighted meta-analysis using the metafor R package (https://cran.r-project.org/web/packages/metafor/).

#### Schizophrenia PRS PheWAS

We conducted PheWASs for both PRS methods in each of the four healthcare systems using all phecodes with sufficient sample size (at least 100 cases). Logistic regressions between schizophrenia PRSs and each phecode were run with 10 ancestry principal components, median age within the medical record calculated for each individual using all of their records in the EHR, sex, genotyping platform, and genotyping batch when available, included as covariates using the PheWAS R package^29^. We used a Bonferroni correction for establishing statistical significance based on the number of phecodes tested at each site. We then meta-analyzed PheWAS effects across healthcare systems within a given PRS method with a fixed-effect inverse variance-weighted model using the PheWAS R package. Phecodes significantly associated with schizophrenia PRS in the PheWAS meta-analysis were carried forward for a follow-up analysis in which we quantified the risk of the phecode at the extremes of the PRS distribution at each site. Effects were combined across sites through meta-analysis using the metafor R package.

#### Sensitivity Analyses to Assess Secondary Effects of Schizophrenia

To explore whether pleiotropic effects of the schizophrenia PRS were mediated by the diagnosis of schizophrenia itself or by the prescription of antipsychotic medications (the most common treatment for schizophrenia), we conducted four follow-up PheWAS analyses. Given the similarity of primary PheWAS results from the two PRS methods, sensitivity analyses were conducted using the LD-pruned PRS method only. For each follow-up analysis, a PheWAS analysis was conducted as above, with only one of the following alterations: an additional covariate for diagnosis of psychotic disorders (phecode 295; the broadest schizophrenia-related phecode), an additional covariate for any prescriptions of antipsychotic medication, removing psychosis cases (phecode 295), and removing patients with any antipsychotic medication prescription history.

## Results

Our sample included 106,160 patients (56% female) across four large US healthcare systems that had collectively received over 35 million ICD-9/10 billing codes. The median length of the electronic health record across sites ranged from 8-15 years and patients had a median range of 52-142 unique visits (Table 1).

**Table 1.**
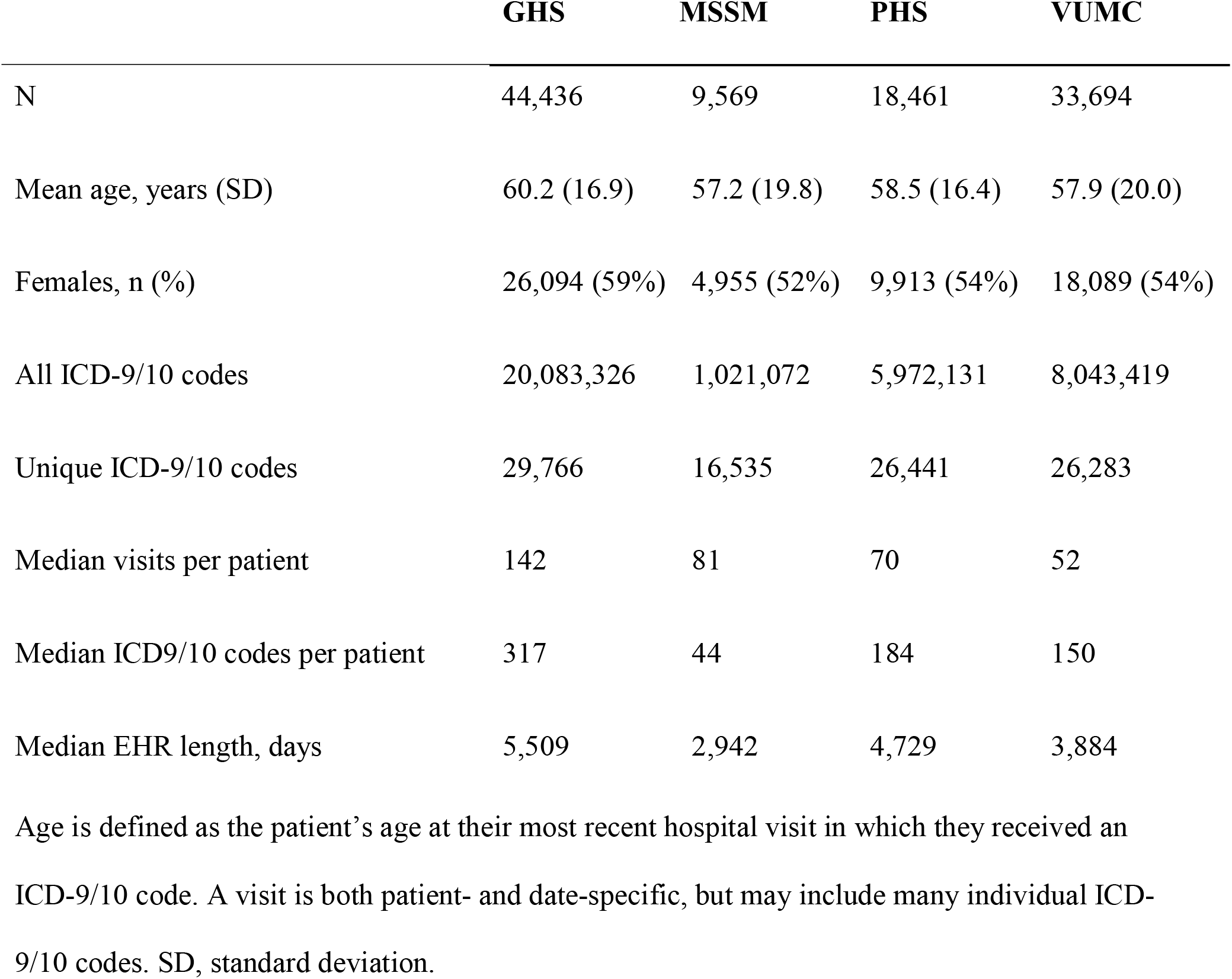
Demographics and Clinical Characteristics

### Penetrance of Schizophrenia PRS in Healthcare Systems

Polygenic risk scores were robustly associated with schizophrenia in the cross-site meta-analysis (OR per standard deviation increase in PRS = 1.55 [95% confidence interval (CI), 1.4-1.7], *p* = 4.48 x 10^-16^) (Table S1); extremely similar effects were observed using the Bayesian PRS (Table S2), as well as in each individual healthcare system (Table S3; Table S4). Absolute risk for schizophrenia in the top decile was 0.8% (Figure 1), equating to 1.9-fold increased odds of schizophrenia compared to those below the 90th percentile (95% CI, 1.5-2.4, *p* = 7.81 x 10^-8^) and 3.3-fold increased odds compared to the bottom decile (95% CI, 2.1-5.2, *p* = 1.16 x 10^-7^) (Figure 2; Table 2). Similarly, for the Bayesian PRS, absolute risk for the top decile was 1.0% (Figure 1), with an OR of 2.3 compared to the bottom 90^th^ percentile (95% CI, 1.9-2.9, *p* = 1.98 x 10^-14^) and 4.6 compared to the bottom decile (95% CI, 2.9-7.3, *p* = 1.37 x 10^-10^) (Figure S1; Table 2).

**Figure 1.**
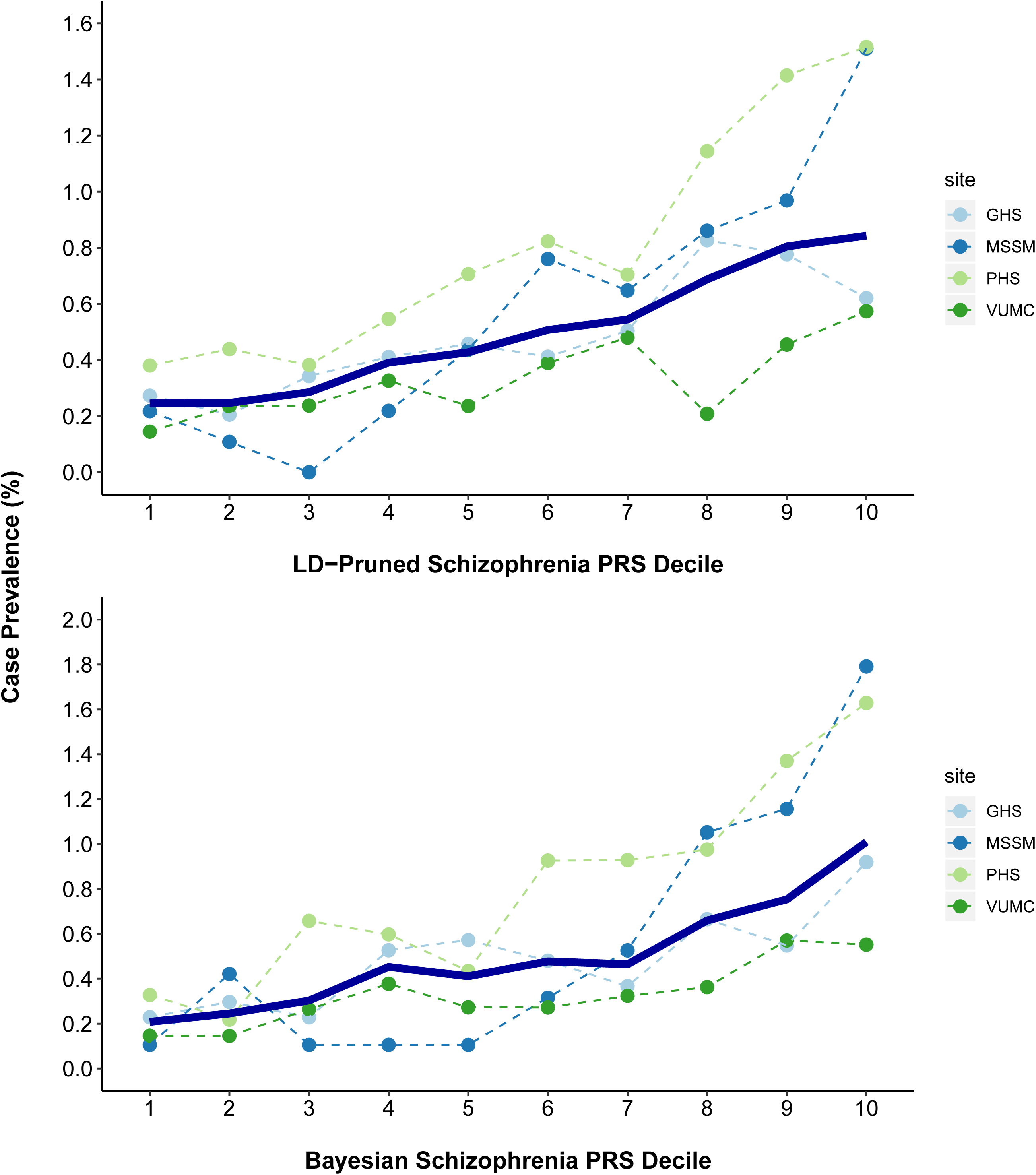
Schizophrenia Case Prevalence by PRS Decile. Schizophrenia case prevalence by site (dashed lines) and across all healthcare systems (solid line) was plotted by schizophrenia PRS decile for both PRS methods. GHS, Geisinger Health System; MSSM, Mount Sinai School of Medicine; PHS, Partners Healthcare System; VUMC, Vanderbilt University Medical Center; PRS, polygenic risk score.

**Figure 2.**
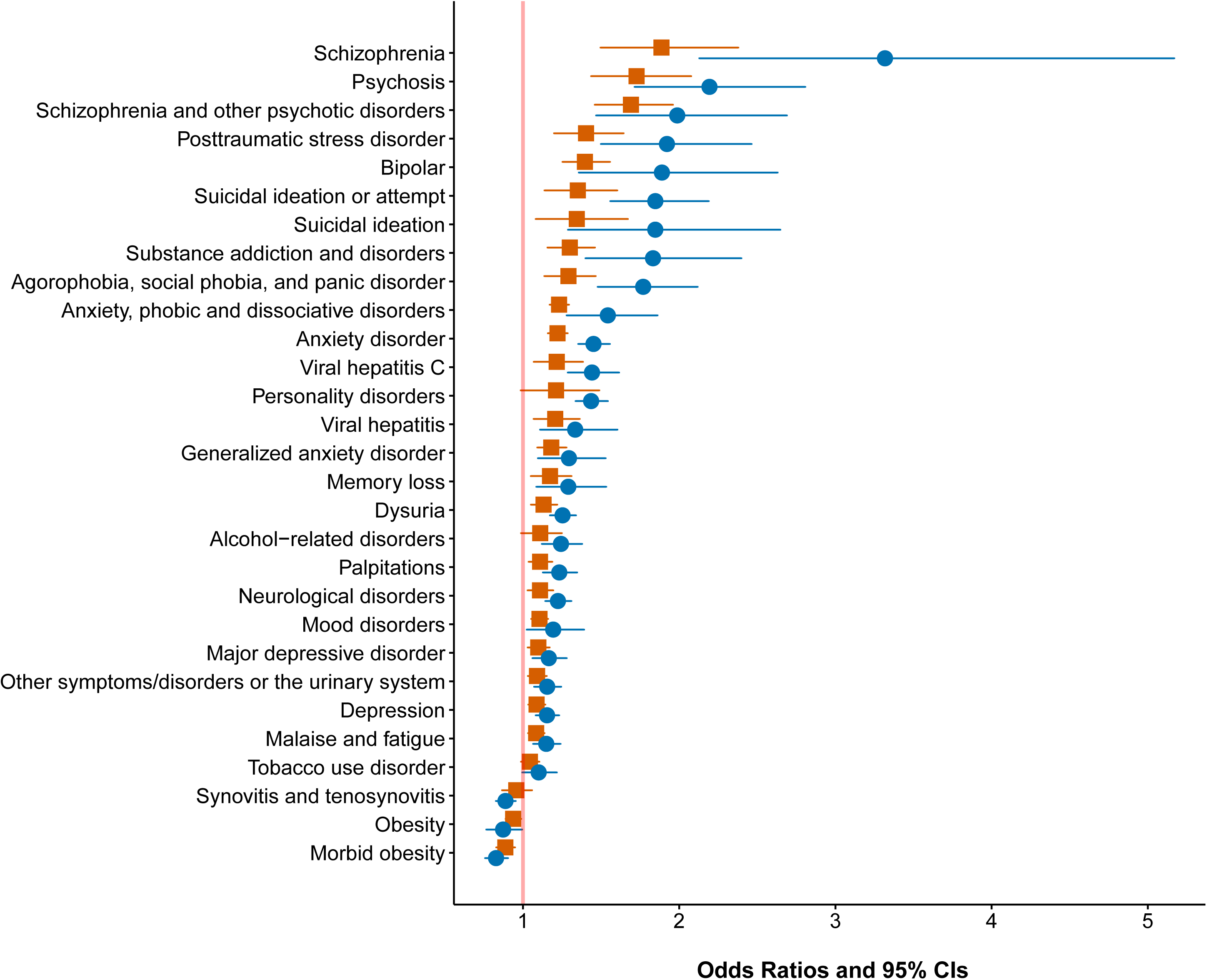
Odds Ratios for Top Schizophrenia PRS Decile. Odds ratios and 95% confidence intervals for phenotypes significant in LD-pruned PRS PheWAS meta-analysis were plotted for the top PRS decile with reference to both the remaining 90% (red squares) and the bottom decile (blue circles). The vertical red line reflects no change in risk (OR = 1).

**Table 2.** Odds Ratios for Schizophrenia and Psychotic Disorders

|  | PRS | Risk | Risk Case | Reference | Reference |  |  |  |
| --- | --- | --- | --- | --- | --- | --- | --- | --- |
|  |  |  |  |  | Case | OR | 95% CI | P Value |
|  | Method | Group | Prevalence | Group | Prevalence |  |  |  |
| **Schizophrenia** | LD-pruned | Top 10% | 0.8% | Remaining 90% | 0.5% | 1.9 | 1.5-2.4 | $7.81 \times 10^{-8}$ |
| | LD-pruned | Top 10% | 0.8% | Bottom 10% | 0.2% | 3.3 | 2.1-5.2 | $1.16 \times 10^{-7}$ |
| | Bayesian | Top 10% | 1.0% | Remaining 90% | 0.4% | 2.3 | 1.9-2.9 | $1.98 \times 10^{-14}$ |
| | Bayesian | Top 10% | 1.0% | Bottom 10% | 0.2% | 4.6 | 2.9-7.3 | $1.37 \times 10^{-10}$ |
| **Schizophrenia and**<br>**Related Psychotic**<br>**Disorders** | LD-pruned | Top 10% | 2.1% | Remaining 90% | 1.3% | 1.7 | 1.5-2.0 | $2.00 \times 10^{-12}$ |
| | LD-pruned | Top 10% | 2.1% | Bottom 10% | 0.9% | 2.2 | 1.7-2.8 | $4.14 \times 10^{-10}$ |
| | Bayesian | Top 10% | 2.1% | Remaining 90% | 1.3% | 1.6 | 1.4-1.9 | $1.75 \times 10^{-10}$ |
| | Bayesian | Top 10% | 2.1% | Bottom 10% | 1.0% | 2.1 | 1.6-2.7 | $2.75 \times 10^{-9}$ |

Overall sample case prevalence was 0.5% for schizophrenia and 1.4% for schizophrenia and related psychotic disorders. PRS, polygenic risk score; OR, odds ratio; CI, confidence interval.

### Schizophrenia PRS PheWAS

After excluding codes for which no site had at least 100 cases, we conducted PheWAS using 1359 disease categories for two PRS methods. The cross-site LD-pruned PRS PheWAS meta-analysis yielded significant associations between schizophrenia PRSs and 29 medical phenotypes including schizophrenia (Table S1; Figure 3). Very similar results were observed using the Bayesian PRS (Table S2) and at each site (Table S3; Table S4). As shown, the strongest cross-site associations were with psychiatric phenotypes for which positive genetic correlations with schizophrenia have been reported, including bipolar disorder, depression, substance use disorders, and anxiety disorders^9^. We additionally found associations with personality disorders, suicidal behavior, neurological disorders, memory loss, viral hepatitis, urinary syndromes and nonspecific somatic symptoms. Obesity and synovitis were inversely associated with schizophrenia PRSs. Effect sizes for all significant phenotypes were plotted in Figures 2 and S2.

**Figure 3.**
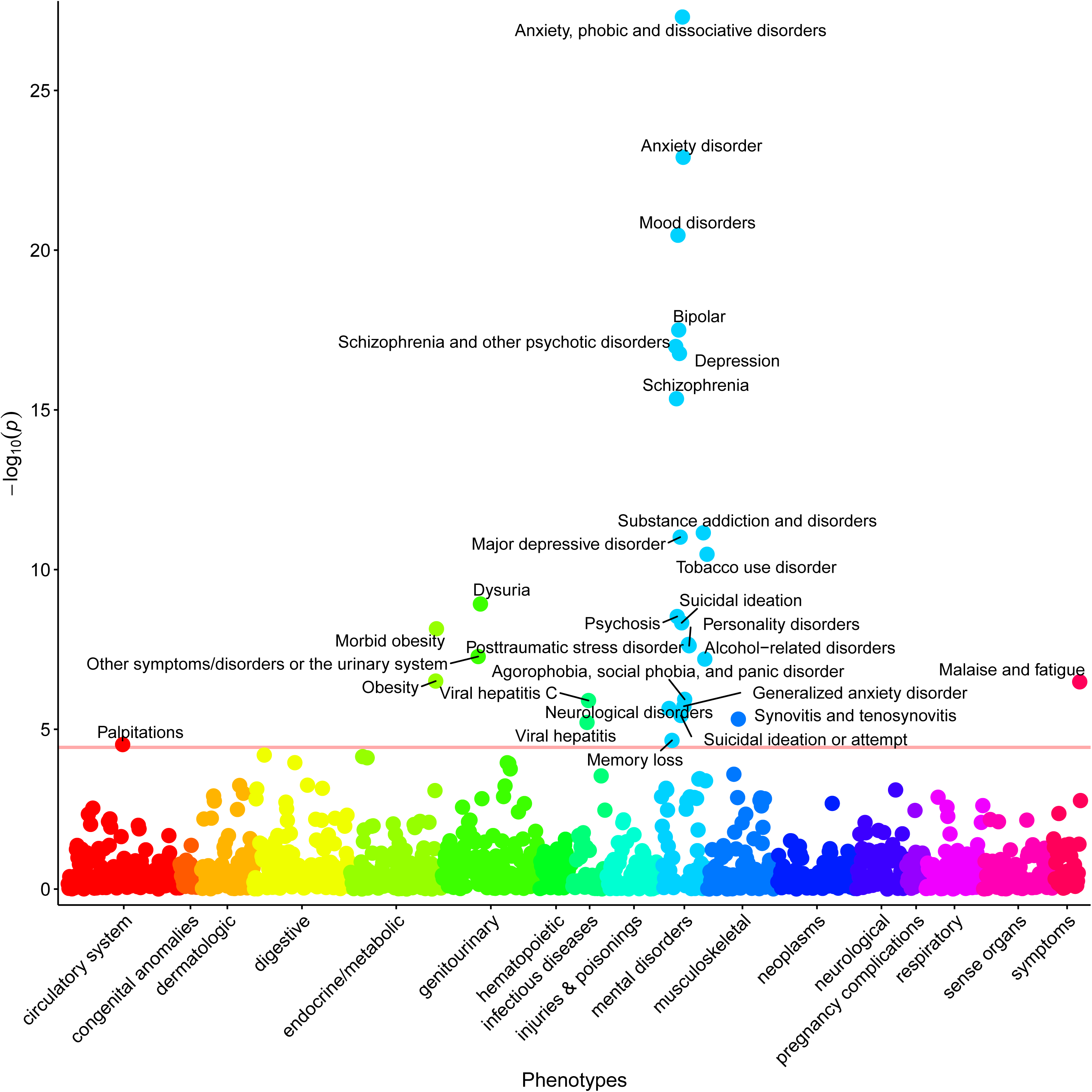
Schizophrenia PRS PheWAS Meta-Analysis. Manhattan plot for phenome-wide association with LD-pruned schizophrenia polygenic risk scores meta-analyzed across four healthcare systems (1359 phenotypes; 106,160 patients). The *x* axis is phenotype (grouped by broad disease category) and the *y* axis is significance (–log_10_ *P*; 2-tailed) of association derived by logistic regression. The red line shows phenome-wide level significance (3.7 x 10^-5^) using Bonferroni correction and all phenotypes passing this threshold are labeled. All significant effects were positive (i.e., higher polygenic risk scores resulted in higher incidence of the phenotype) with three exceptions: morbid obesity, obesity, and synovitis and tenosynovitis.

### Sensitivity Analyses to Assess Secondary Effects of Schizophrenia

We explored whether some of the observed associations might be mediated through a clinical diagnosis of schizophrenia or antipsychotic medication use through a series of sensitivity analyses. Nearly all associations remained significant across all sensitivity analyses (Table S5; Figure S2), although for some phecodes, there was minor variability. Nonetheless, in every analysis, phecodes related to anxiety disorders, mood disorders, substance use disorders, obesity phenotypes, urinary syndromes, and malaise and fatigue remained significant. Associations with suicidal behavior, personality disorders, neurological disorders, memory loss, synovitis and tenosynovitis, and viral hepatitis were less robust, although they remained top phenotypes consistently (Table S5).

## Discussion

We investigated the impact of genetic risk for schizophrenia across the medical phenome in 106,160 patients from four large healthcare systems. Several findings from our analyses are particularly noteworthy. First, externally-derived polygenic risk scores for schizophrenia robustly detected risk for diagnosis of schizophrenia in real-world healthcare settings (*p*’s < 4.48 x 10^-16^). The effect sizes (Table 2) were similar to those observed for corresponding PRSs for atrial fibrillation, type 2 diabetes, inflammatory bowel disease, many common cancers^13, 31^. Second, we leveraged the phenome-wide data available in EHRs to conduct the first psychiatric PRS PheWAS in multiple U.S. healthcare systems, revealing a range of pleiotropic relationships.

While we reported strong associations with schizophrenia, the effect sizes were more modest than those reported in schizophrenia case-control cohorts ascertained for research purposes. For example, in the original report by the PGC from which the risk scores were derived, individuals in the top decile of schizophrenia PRS relative to the bottom decile had a 7.8-20.3 increased odds of schizophrenia^14^, whereas we observed odds ratios of 3.3 and 4.6, depending on the PRS method (Table 2). There are several potential reasons for this discrepancy. First, cases in the PGC meta-analysis met relatively stringent criteria based on clinical interviews by trained research personnel, and control ascertainment often included screening for history of psychiatric or neurological disorders. This approach, typical for research samples, maximizes power for genetic discovery by extreme sampling from the tails of the genetic liability distribution. In contrast, our analysis was expressly designed to approximate use of a PRS in a typical clinical setting by applying a simple definition for both cases (two or more schizophrenia-related codes) and controls (no schizophrenia-related codes). Thus, although the effect size we observed is likely attenuated due to some degree of misclassification, it may better reflect results that would be seen in real-world clinical settings where PRSs are applied to a broad healthcare population with little *a priori* knowledge of clinical symptoms. In addition, we did not restrict the age range of cases and controls, which may have further reduced the apparent effect size of the schizophrenia PRS (some individuals in our sample who have not yet reached the age of illness onset may have been misclassified as controls).

Although the PRS effects we observed were not large enough on their own to stratify risk in a clinical setting (i.e., to discriminate between cases and controls on an individual level with high accuracy), they are comparable to those of risk factors in established risk calculators. For example, two well-established coronary artery disease (CAD) risk factors – smoking and diabetes – were estimated in the Framingham Heart Study to have hazard ratios ≤ 2.0^32^ – similar to the observed risk for the top schizophrenia PRS decile here. Additionally, in a risk calculator for the transition to psychosis among high-risk individuals – one of the few individualized risk calculators developed within psychiatry – the best predictor was a symptom severity index with a hazard ratio of 2.1 (95% CI, 1.6-2.7)^33^. While this risk calculator was not validated for clinical use, it does reflect effects of variables used by clinicians to assess risk in the clinic.

In light of this, we speculate that incorporating genetic risk could be impactful within psychiatry, especially as enhanced performance may be possible through a variety of means. For example, we implemented two PRS methods, a standard LD-pruning approach and a newer, Bayesian one, to evaluate the robustness and consistency of our results. While the differences in results were not large, the Bayesian method produced larger effect estimates overall, including for schizophrenia (Table 2). These findings support the use of newer risk scoring methods that can incorporate more genetic variants by directly modeling LD structure. Precision of PRSs may also increase through larger discovery sample sizes^12^ and with refinement of EHR-based case definitions. Nonetheless, it remains to be seen whether combining PRS risk estimates with other clinical predictors can meaningfully contribute to individualized risk assessment in psychiatry.

Schizophrenia PRSs were also associated with broader effects on mental health including increased risk for anxiety, mood, substance use, personality, and neurological disorders, as well as memory loss and suicidal behavior. Anxiety, mood, and substance use disorders have all previously been linked to genetic risk for schizophrenia^9, 34–36^ and our results confirm in a clinical setting that these disorders share genetic risk. Certain personality disorders have also been linked to genetic liability for schizophrenia^37, 38^ (e.g., schizotypal or schizoid) and there is some evidence that personality dimensions in adolescence predict future psychopathology, including schizophrenia^39^. Similarly, family history of schizophrenia has been associated with suicidal behavior^40^. However, results from our sensitivity analyses suggested that the relationships between schizophrenia and neurological disorders, personality disorders, suicidal behavior, and memory loss may be consequences of a schizophrenia diagnosis rather than due to shared genetic risk (Figure S2).

Genetic liability for schizophrenia was associated with many non-psychiatric syndromes as well, including obesity, urinary syndromes, viral hepatitis, synovitis and tenosynovitis, and malaise and fatigue. Intriguingly, obesity and morbid obesity were significantly negatively associated with schizophrenia PRSs (Table S1). This is somewhat surprising given the known phenotypic correlation between schizophrenia and obesity^41^. Nonetheless, three prior reports found significant inverse genetic correlations between body mass index and schizophrenia^42–44^, while a fourth reported an inverse, but non-significant relationship^45^. This may suggest that increased rates of obesity among patients with schizophrenia may be a consequence of the disease, potentially due to antipsychotic use or poor support for proper nutrition. We also found an inverse association between genetic liability for schizophrenia and diabetes, but only in sensitivity analyses controlling for a schizophrenia diagnosis or antipsychotic medication history. It may be that this negative genetic correlation was attenuated in the primary analysis (i.e., including patients with schizophrenia and antipsychotic medication history with no statistical control) due to diabetes-promoting effects of antipsychotic medications within the same individuals that were at high genetic risk for schizophrenia^41^. In general, pleiotropic effects may have implications for risk communication if PRS testing is deployed in clinical settings in the future.

Our results should be interpreted in light of several limitations. First, due to small numbers of patients of other ancestries, our analyses were restricted to patients of European descent, and the generalizability to individuals of non-European ancestry remains to be determined. Second, our phenotype definitions relied on very simple rules and disregarded many variables of potential importance including medical history of related disorders, setting of diagnosis (i.e., in- or outpatient; physician specialty), and treatment for the disease of interest. This was by design in order to mimic a real-world clinical population where PRSs may be implemented for clinical decision support, however, the approach is sensitive to misclassifications that occur in a clinical setting. Future work refining case and control definitions using natural language processing algorithms may improve the predictive performance of PRSs and other risk factors for clinically-derived phenotypes^47, 48^. Third, our results varied to some degree between sites (Table S3; Table S4), perhaps most notably for schizophrenia, suggesting that demographic and disease distributions in any given healthcare system will influence penetrance and pleiotropy. However, we tested for between-site heterogeneity for schizophrenia, and though this test has relatively low power, it showed no evidence of significant heterogeneity (*p*’s > .45). Relatedly, disease prevalence was often lower in the overall healthcare system relative to the participants enrolled in the biobanks (a subset of those patients) (Table S6). In general, case prevalence in the biobanks was more representative of population-level prevalence than was that in the healthcare systems, suggesting that the discrepancies may be due to biobank patients generally having a longer duration of EHR follow-up and therefore more opportunity to receive a diagnosis than patients in the overall healthcare system (Table S6). Finally, although our analyses comprise the largest test of a schizophrenia PRS in EHR data to date, additional phenotypes may show significant association in future, larger-scale PheWAS.

In conclusion, we demonstrate that an available measure of polygenic risk for schizophrenia is robustly associated with schizophrenia across four large healthcare systems using EHR data. While the observed penetrance of schizophrenia PRS is attenuated in these settings compared to prior estimates derived from research cohorts, effect sizes are comparable to those seen for risk factors commonly used in clinical settings. We also find that polygenic risk for schizophrenia has pleiotropic effects on related psychiatric disorders as well as several non-psychiatric symptoms and syndromes. Our results provide an initial indication of the opportunities and limitations that may arise with the future application of PRS testing in healthcare systems.

## Disclosures and acknowledgements

Dr. Smoller is an unpaid member of the Bipolar/Depression Research Community Advisory Panel of 23andMe. Dr. Kirchner received funding from Regeneron Genetics Center as part of the DiscovEHR study. All other authors have nothing to report.

This work was supported in part by NIMH grant R01MH118233 (J.W.S. and L.K.D.) We would also like to acknowledge the Partners Biobank for providing samples, genomic data, and health information data. J.W.S. was supported in part by an NHGRI grant supporting the eMERGE Network (U01HG008685) and T.G. was supported in part by an NIA grant (K99AG054573). J.W.S. is also a Tepper Family MGH Research Scholar and supported in part by the Demarest Lloyd, Jr. Foundation.

The VUMC components of the project were conducted in part using the resources of the Advanced Computing Center for Research and Education at Vanderbilt University, Nashville, TN. The datasets used for the project were obtained from VUMCs Synthetic Derivative and BioVU, which are supported by numerous sources: institutional funding, private agencies, and federal grants. These include the NIH funded Shared Instrumentation Grant S10RR025141; and CTSA grants UL1TR002243, UL1TR000445, and UL1RR024975 from NCATS/NIH. Additional funding provided by the NIH through grants P50GM115305 and U19HL065962. The authors wish to acknowledge the expert technical support of the VANTAGE and VANGARD core facilities, supported in part by the Vanderbilt-Ingram Cancer Center (P30 CA068485) and Vanderbilt Vision Center (P30 EY08126). Additionally, J.D. was supported by the Canadian Institutes of Health Research (MFE-142936), L.K.D. was supported by R01MH113362, and D.R. was supported by R01MH111776.

The authors would also like to thank the MyCode Community Health Initiative participants for their permission to utilize their health and genomics information in the DiscovEHR collaboration. The DiscovEHR study was funded in part by the Regeneron Genetics Center and the Geisinger Clinic was funded by grant U01HG008679.

Finally, we would like to acknowledge the the Andrea and Charles Bronfman Philanthropies for supporting the Mount Sinai BioMe Biobank. E.A.S. was supported in part by several NIMH grants (U01MH109536, R01MH106531, R01MH095034).

## References

1. Walker ER, McGee RE, Druss BG. Mortality in mental disorders and global disease burden implications a systematic review and meta-analysis. JAMA Psychiatry. 2015. doi:10.1001/jamapsychiatry.2014.2502.

2. Vos T, Abajobir AA, Abbafati C, et al. Global, regional, and national incidence, prevalence, and years lived with disability for 328 diseases and injuries for 195 countries, 1990-2016: A systematic analysis for the Global Burden of Disease Study 2016. Lancet. 2017. doi:10.1016/S0140-6736(17)32154-2.

3. Steel Z, Marnane C, Iranpour C, et al. The global prevalence of common mental disorders: A systematic review and meta-analysis 1980-2013. Int J Epidemiol. 2014. doi:10.1093/ije/dyu038.

4. Ahrnsbrak R, Bose J, Hedden SL, Lipari RN, Park-Lee E. Key substance use and mental health indicators in the United States: results from the 2016 national survey on drug use and health. Subst Abus Ment Heal Serv Adm. 2016. doi:10.1016/j.drugalcdep.2016.10.042.

5. Albert N, Melau M, Jensen H, Hastrup LH, Hjorthøj C, Nordentoft M. The effect of duration of untreated psychosis and treatment delay on the outcomes of prolonged early intervention in psychotic disorders. npj Schizophr. 2017. doi:10.1038/s41537-017-0034-4.

6. Tang JYM, Chang WC, Hui CLM, et al. Prospective relationship between duration of untreated psychosis and 13-year clinical outcome: A first-episode psychosis study. Schizophr Res. 2014. doi:10.1016/j.schres.2014.01.022.

7. Amminger GP, Edwards J, Brewer WJ, Harrigan S, McGorry PD. Duration of untreated psychosis and cognitive deterioration in first-episode schizophrenia. Schizophr Res. 2002. doi:10.1016/S0920-9964(01)00278-X.

8. Wang PS, Berglund P, Olfson M, Pincus HA, Wells KB, Kessler RC. Failure and delay in initial treatment contact after first onset of mental disorders in the National Comorbidity Survey Replication. Arch Gen Psychiatry. 2005. doi:10.1001/archpsyc.62.6.603.

9. Brainstorm Consortium TB, Anttila V, Bulik-Sullivan B, et al. Analysis of shared heritability in common disorders of the brain. Science. 2018;360(6395):eaap8757. doi:10.1126/science.aap8757.

10. Smoller JW, Andreassen OA, Edenberg HJ, Faraone S V., Glatt SJ, Kendler KS. Psychiatric genetics and the structure of psychopathology. Mol Psychiatry. January 2018:1. doi:10.1038/s41380-017-0010-4.

11. Vassos E, Di Forti M, Coleman J, et al. An Examination of Polygenic Score Risk Prediction in Individuals With First-Episode Psychosis. Biol Psychiatry. 2017. doi:10.1016/j.biopsych.2016.06.028.

12. Zheutlin AB, Ross DA. Polygenic Risk Scores: What Are They Good For? Biol Psychiatry. 2018. doi:10.1016/j.biopsych.2018.04.007.

13. Khera A V., Chaffin M, Aragam KG, et al. Genome-wide polygenic scores for common diseases identify individuals with risk equivalent to monogenic mutations. Nat Genet. August 2018:1. doi:10.1038/s41588-018-0183-z.

14. Schizophrenia Working Group of the Psychiatric Genomics C. Biological insights from 108 schizophrenia-associated genetic loci. Nature. 2014;511(7510):421–427. doi:10.1038/nature13595.

15. Crawford DC, Crosslin DR, Tromp G, et al. EMERGEing progress in genomics-the first seven years. Front Genet. 2014. doi:10.3389/fgene.2014.00184.

16. Gottesman O, Kuivaniemi H, Tromp G, et al. The Electronic Medical Records and Genomics (eMERGE) Network: Past, present, and future. Genet Med. 2013. doi:10.1038/gim.2013.72.

17. Smoller JW. The use of electronic health records for psychiatric phenotyping and genomics. American Journal of Medical Genetics, Part B: Neuropsychiatric Genetics. 2017.

18. Solovieff N, Cotsapas C, Lee PH, Purcell SM, Smoller JW. Pleiotropy in complex traits: Challenges and strategies. Nat Rev Genet. 2013. doi:10.1038/nrg3461.

19. Carey DJ, Fetterolf SN, Davis FD, et al. The Geisinger MyCode community health initiative: An electronic health record-linked biobank for precision medicine research. Genet Med. 2016. doi:10.1038/gim.2015.187.

20. Belbin GM, Odgis J, Sorokin EP, et al. Genetic identification of a common collagen disease in puerto ricans via identity-by-descent mapping in a health system. Elife. 2017. doi:10.7554/eLife.25060.

21. Karlson EW, Boutin NT, Hoffnagle AG, Allen NL. Building the partners healthcare biobank at partners personalized medicine: Informed consent, return of research results, recruitment lessons and operational considerations. J Pers Med. 2016. doi:10.3390/jpm6010002.

22. Danciu I, Cowan JD, Basford M, et al. Secondary use of clinical data: The Vanderbilt approach. J Biomed Inform. 2014. doi:10.1016/j.jbi.2014.02.003.

23. Duncan L, Shen H, Gelaye B, et al. Analysis of Polygenic Score Usage and Performance in Diverse Human Populations. bioRxiv. 2018. doi:10.1101/398396.

24. Martin AR, Kanai M, Kamatani Y, Okada Y, Neale BM, Daly MJ. Current clinical use of polygenic scores will risk exacerbating health disparities. bioRxiv. 2019. doi:10.1101/441261.

25. Euesden J, Lewis CM, O’Reilly PF. PRSice: Polygenic Risk Score software. Bioinformatics. 2015. doi:10.1093/bioinformatics/btu848.

26. Ge T, Chen C-Y, Ni Y, Feng Y-CA, Smoller JW. Polygenic Prediction via Bayesian Regression and Continuous Shrinkage Priors. bioRxiv. 2018. doi:10.1101/416859.

27. Chang CC, Chow CC, Tellier LCAM, Vattikuti S, Purcell SM, Lee JJ. Second-generation PLINK: Rising to the challenge of larger and richer datasets. Gigascience. 2015. doi:10.1186/s13742-015-0047-8.

28. Wei WQ, Bastarache LA, Carroll RJ, et al. Evaluating phecodes, clinical classification software, and ICD-9-CM codes for phenome-wide association studies in the electronic health record. PLoS One. 2017. doi:10.1371/journal.pone.0175508.

29. Carroll RJ, Bastarache L, Denny JC. R PheWAS: Data analysis and plotting tools for phenome-wide association studies in the R environment. Bioinformatics. 2014. doi:10.1093/bioinformatics/btu197.

30. Wei WQ, Teixeira PL, Mo H, Cronin RM, Warner JL, Denny JC. Combining billing codes, clinical notes, and medications from electronic health records provides superior phenotyping performance. J Am Med Informatics Assoc. 2016. doi:10.1093/jamia/ocv130.

31. Fritsche LG, Gruber SB, Wu Z, et al. Association of Polygenic Risk Scores for Multiple Cancers in a Phenome-wide Study: Results from The Michigan Genomics Initiative. American Journal of Human Genetics. 2018.

32. D’Agostino RB, Vasan RS, Pencina MJ, et al. General cardiovascular risk profile for use in primary care: The Framingham heart study. Circulation. 2008. doi:10.1161/CIRCULATIONAHA.107.699579.

33. Cannon TD, Yu C, Addington J, et al. An individualized risk calculator for research in prodromal psychosis. Am J Psychiatry. 2016;173(10):980–988. doi:10.1176/appi.ajp.2016.15070890.

34. Gandal MJ, Haney JR, Parikshak NN, et al. Shared molecular neuropathology across major psychiatric disorders parallels polygenic overlap. Science. 2018;359(6376):693–697. doi:10.1126/science.aad6469.

35. Smoller JW, Craddock N, Kendler K, et al. Identification of risk loci with shared effects on five major psychiatric disorders: a genome-wide analysis. Lancet. 2013;381(9875):1371–1379. doi:10.1016/S0140-6736(12)62129-1.

36. Mistry S, Harrison JR, Smith DJ, Escott-Price V, Zammit S. The use of polygenic risk scores to identify phenotypes associated with genetic risk of schizophrenia: Systematic review. Schizophr Res. 2017. doi:10.1016/j.schres.2017.10.037.

37. Nelson MT, Seal ML, Pantelis C, Phillips LJ. Evidence of a dimensional relationship between schizotypy and schizophrenia: A systematic review. Neurosci Biobehav Rev. 2013. doi:10.1016/j.neubiorev.2013.01.004.

38. Bigdeli TB, Bacanu SA, Webb BT, et al. Molecular validation of the schizophrenia spectrum. Schizophr Bull. 2014. doi:10.1093/schbul/sbt122.

39. Newton-Howes G, Horwood J, Mulder R. Personality characteristics in childhood and outcomes in adulthood: Findings from a 30 year longitudinal study. Aust N Z J Psychiatry. 2015. doi:10.1177/0004867415569796.

40. Laursen TM, Trabjerg BB, Mors O, et al. Association of the polygenic risk score for schizophrenia with mortality and suicidal behavior - A Danish population-based study. Schizophr Res. 2017. doi:10.1016/j.schres.2016.12.001.

41. Annamalai A, Kosir U, Tek C. Prevalence of obesity and diabetes in patients with schizophrenia. World J Diabetes. 2017. doi:10.4239/wjd.v8.i8.390.

42. So H-C, Chau K-L, Ao F-K, Mo C-H, Sham P-C. Exploring shared genetic bases and causal relationships of schizophrenia and bipolar disorder with 28 cardiovascular and metabolic traits. Psychol Med. 2018. doi:10.1017/S0033291718001812.

43. Akiyama M, Okada Y, Kanai M, et al. Genome-wide association study identifies 112 new loci for body mass index in the Japanese population. Nat Genet. 2017. doi:10.1038/ng.3951.

44. Duncan LE, Shen H, Ballon JS, Hardy K V, Noordsy DL, Levinson DF. Genetic Correlation Profile of Schizophrenia Mirrors Epidemiological Results and Suggests Link Between Polygenic and Rare Variant (22q11.2) Cases of Schizophrenia. Schizophr Bull. 2017. doi:10.1093/schbul/sbx174.

45. Bulik-Sullivan B, Finucane HK, Anttila V, et al. An atlas of genetic correlations across human diseases and traits. Nat Genet. 2015. doi:10.1038/ng.3406.

46. Carson CM, Phillip N, Miller BJ. Urinary tract infections in children and adolescents with acute psychosis. Schizophr Res. 2017. doi:10.1016/j.schres.2016.11.004.

47. Perlis RH, Iosifescu D V., Castro VM, et al. Using electronic medical records to enable large-scale studies in psychiatry: Treatment resistant depression as a model. Psychol Med. 2012. doi:10.1017/S0033291711000997.

48. Castro VM, Minnier J, Murphy SN, et al. Validation of electronic health record phenotyping of bipolar disorder cases and controls. Am J Psychiatry. 2015. doi:10.1176/appi.ajp.2014.14030423.

