## Supplemental Methods and Supplemental Table and Figure Legends for "Penetrance and pleiotropy of polygenic risk scores for schizophrenia in 106,160 patients across four healthcare systems"

Supplementary Methods

Supplementary Tables (see SupplementaryTables.xlsx)

Table S1. LD-pruned Schizophrenia PRS PheWAS Meta-Analysis

Table S2. Bayesian Schizophrenia PRS PheWAS Meta-Analysis

Table S3. LD-pruned Schizophrenia PRS PheWAS Results by Site

Table S4. Bayesian Schizophrenia PRS PheWAS Results by Site

Table S5. Sensitivity Analyses for Schizophrenia PRS PheWAS

Table S6. Biobank and Hospital Demographics and Case Prevalence

Supplementary Figures (see SupplementaryFigures.pdf)

Figure S1. Odds Ratios for Top Bayesian PRS Decile

Figure S2. Sensitivity Analyses for Schizophrenia PRS PheWAS

**Supplementary Methods**

An overview of the genotype quality control methods employed across sites was described in the main text (Methods: Quality Control of Genetic Data). Details for each site were listed below.

*GHS:* The Geisinger patient samples (N = 59,085) were genotyped on the Illumina Human OmniExpress Exome BeadChip (958,497 markers) by the Regeneron Genetics Center. Samples were genotyped in batches of ~20,000-25,000 samples. Individuals and SNPs were filtered based on call rate (<1%), minor allele frequency (<1%), and Hardy-Weinberg equilibrium (HWE) (*p* < 10^-7^). A random individual from pairs of related individuals was removed (pihat > 0.125). Patients with European ancestry determined genetically were retained. Strand alignment, checks, and phasing were performed with SHAPEIT v2; palindromic SNPs were removed during strand alignment. Imputation was performed with IMPUTE2 separately by batch on autosomal chromosomes with the 1000 Genomes Phase 1 reference panel. IMPUTE2 files were converted to hard-call PLINK format using filters for genotype probability (>0.9) and INFO score (>0.7). Batches were merged after imputation in PLINK v1.90. Principal components to include as ancestry covariates were generated in EIGENSOFT.

*MSSM*: A subset of Bio*Me* biobank participants were genotyped on the Illumina Global Screening Array. Samples were blacklisted following genotyping for gender discordance (genotype sex-check results with homozygosity rates more than 0.2 but less than 0.8 were excluded), low sequencing coverage, heterozygosity rates (samples falling outside of three standard deviations from the mean were excluded), contamination, low call rate, and the discovery of duplicates. A random individual from all pairs with apparent relatedness (coefficient > 0.0445) was excluded. Variants were removed for excessive missingness (>3%). SHAPEIT / IMPUTE2 were used to pre-phase and impute genotypes using the 1000 Genomes Phase 3 reference panel. Dosage data was converted to hard genotype calls, and variants with certainty less than 0.9 or INFO < 0.9 were excluded. Self-reported ancestry information was then used to extract the subset of European individuals for the present analysis (N = 9,845).

*PHS:* 25,540 individuals were genotyped on one of three different Illumina arrays (MEGA, MEGA^EX^, and MEG BeadChip). Individuals and SNPs were excluded from each array for call rate (<2%), sex errors, and heterozygosity (|F_het_| > 0.2); then all three arrays were merged using only variants present in all three datasets. SNPs that had missingness rates that differed between chips (>1%) or showed significant array effects (*p* < 10^-6^) in the merged dataset were excluded. To identify and extract individuals of European-American ancestry, we pruned the initial merged dataset by linkage disequilibrium and randomly removed one individual from pairs with apparent relatedness (pihat > 0.2). We merged this dataset with the 1000 Genomes sample^2^ and generated principal components (PCs). 1000 Genomes has labeled individuals within their dataset as belonging to one of five ancestrally distinct super-populations. We used the first four PCs to identify individuals from PHS that clustered with the European 1000 Genomes super-population (N = 19,136). Next, we used 1000 Genomes Phase III reference panel to impute the initial merged dataset (N_SNP_ = 1,345,786) with only unrelated European-American individuals (N = 19,136). We used Eagle v2.3.5 for prephasing and Minimac3 for imputation. Dosage files were converted to hard-call genotypes with PLINK v1.90 using filters for genotyping probability (*P* >.8), INFO score (>.9), and SNP missingness (<2%). After a final round of quality control using filters for violations of HWE (*p* < 10^-4^) and minor allele frequency (<1%), our dataset included 19,136 European-American patients and 6,237,592 variants.

*VUMC:* A subset of BioVU patients (N = 24,262) were genotyped on the Illumina MEGA^EX^ platform of more than 2 million markers. Ruderfer and colleagues described the first phase of quality control elsewhere^1^, including filters for SNP and individual call rate (<2%), minor allele frequency (<1%), violations of HWE (*p* < 5 x 10^-5^), batch effects (*p* < 5 x 10^-5^), heterozygosity (|F_het_| > 0.2), and relatedness (pihat > 0.2). Ancestry principal components were used to identify individuals of European ancestry, and SHAPEIT / IMPUTE2 were used to pre-phase and impute genotypes according to the 1000 Genomes Phase I reference panel. In the second phase of quality control, we converted dosage data to hard genotype calls and excluded variants with certainty less than 0.9 or INFO < 0.95.  After these quality control measures, 18,385 individuals remained. In all subsequent analyses, we used genotype batch and the first 10 ancestry principal components calculated by multidimensional scaling in PLINK v1.90 as covariates.

Supplementary Tables

*See SupplementaryTables.xlxs*

Table S1.

**LD-pruned Schizophrenia PRS PheWAS Meta-Analysis.** Effects surpassing the Bonferroni significance threshold (3.7 x 10^-5^) were highlighted in blue.

Table S2.

**Bayesian Schizophrenia PRS PheWAS Meta-Analysis.** Effects surpassing the Bonferroni significance threshold (3.7 x 10^-5^) were highlighted in red.

Table S3.

**LD-pruned Schizophrenia PRS PheWAS Results by Site.** Phenotypes were listed in rank order of significance from the LD-pruned PRS PheWAS meta-analysis. Individual effects were listed for each site and those that surpassed the site-specific Bonferroni-corrected significance threshold were highlighted in blue (GHS, 1223 phecodes, *p* < 4.09 x 10^-5^; MSSM, 314 phecodes, *p* < 1.59 x 10^-4^; PHS, 967 phecodes, *p* < 5.17 x 10^-5^; VUMC, 1133 phecodes, *p* < 4.41 x 10^-5^).

Table S4.

**Bayesian Schizophrenia PRS PheWAS Results by Site.** Phenotypes were listed in rank order of significance from the Bayesian PRS PheWAS meta-analysis. Individual effects were listed for each site and those that surpassed the site-specific Bonferroni-corrected significance threshold were highlighted in red (GHS, 1223 phecodes, *p* < 4.09 x 10^-5^; MSSM, 314 phecodes, *p* < 1.59 x 10^-4^; PHS, 967 phecodes, *p* < 5.17 x 10^-5^; VUMC, 1133 phecodes, *p* < 4.41 x 10^-5^).

Table S5.

**Sensitivity Analyses for Schizophrenia PRS PheWAS.** Phenotypes were listed in rank order of significance from the LD-pruned schizophrenia PRS PheWAS meta-analysis. Effects that surpassed the Bonferroni-corrected significance threshold (3.7 x 10^-5^) were highlighted in blue.

Table S6.

**Biobank and Hospital Demographics and Case Prevalence.** Age is defined as the patient’s age at their most recent hospital visit in which they received an ICD-9/10 code. Hospital demographics and prevalence were calculated using patients with three or more EHR visits, one of which at age 10 or older.

Supplementary Figures

*See SupplementaryFigures.pdf*

Figure S1.

**Odds Ratios for Top Bayesian PRS Decile.** Odds ratios and 95% confidence intervals for phenotypes significant in Bayesian PRS meta-analysis were plotted for the top PRS decile with reference to both the remaining 90% (red squares) and the bottom decile (blue circles). The vertical red line reflects no change in risk (OR = 1).

Figure S2.

**Sensitivity Analyses for Schizophrenia PRS PheWAS.** Manhattan plots for sensitivity analyses conducted as follow-up to the primary phenome-wide association analysis with LD-pruned polygenic risk scores. All four meta-analyses use the same methodology as the primary PheWAS with a single alteration – a covariate for phecode 295, the broadest phecode for psychosis (top left); phecode 295 cases excluded (top right); a covariate for any history of antipsychotic medication (bottom left); patients with any antipsychotic medication history excluded (bottom right). For all plots, the *x* axis is phenotype (grouped by broad disease category) and the *y* axis is significance (–log_10_ *P*; 2-tailed) of association derived by logistic regression. The red line shows phenome-wide level significance (3.7 x 10^-5^). All significant effects were positive (i.e., higher polygenic risk scores resulted in higher incidence of the phenotype) with five exceptions: type 2 diabetes, diabetes mellitus, synovitis and tenosynovitis, obesity, and morbid obesity.

**References**

1.	Ruderfer DM, Walsh CG, Aguirre MW, et al. Significant shared heritability underlies suicide attempt and clinically predicted probability of attempting suicide. *bioRxiv*. 2018. doi:10.1101/266411.

2.	The 1000 Genomes Project Consortium. A global reference for human genetic variation. *Nature*. 2015;526(7571):68-74. doi:10.1038/nature15393.
