## Supplementary figures and images for "Penetrance and pleiotropy of polygenic risk scores for schizophrenia in 106,160 patients across four healthcare systems"

### Supplemental Figures

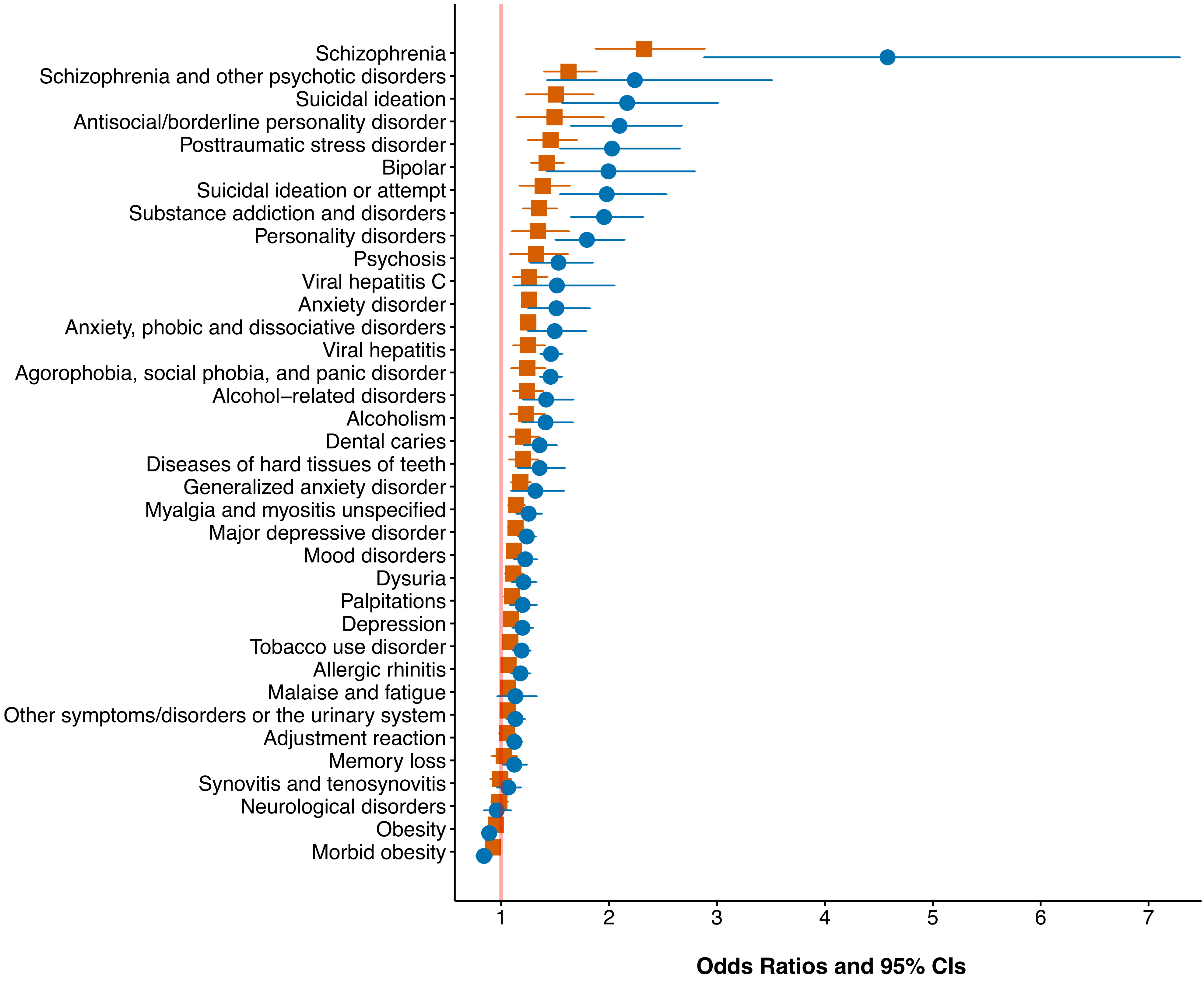

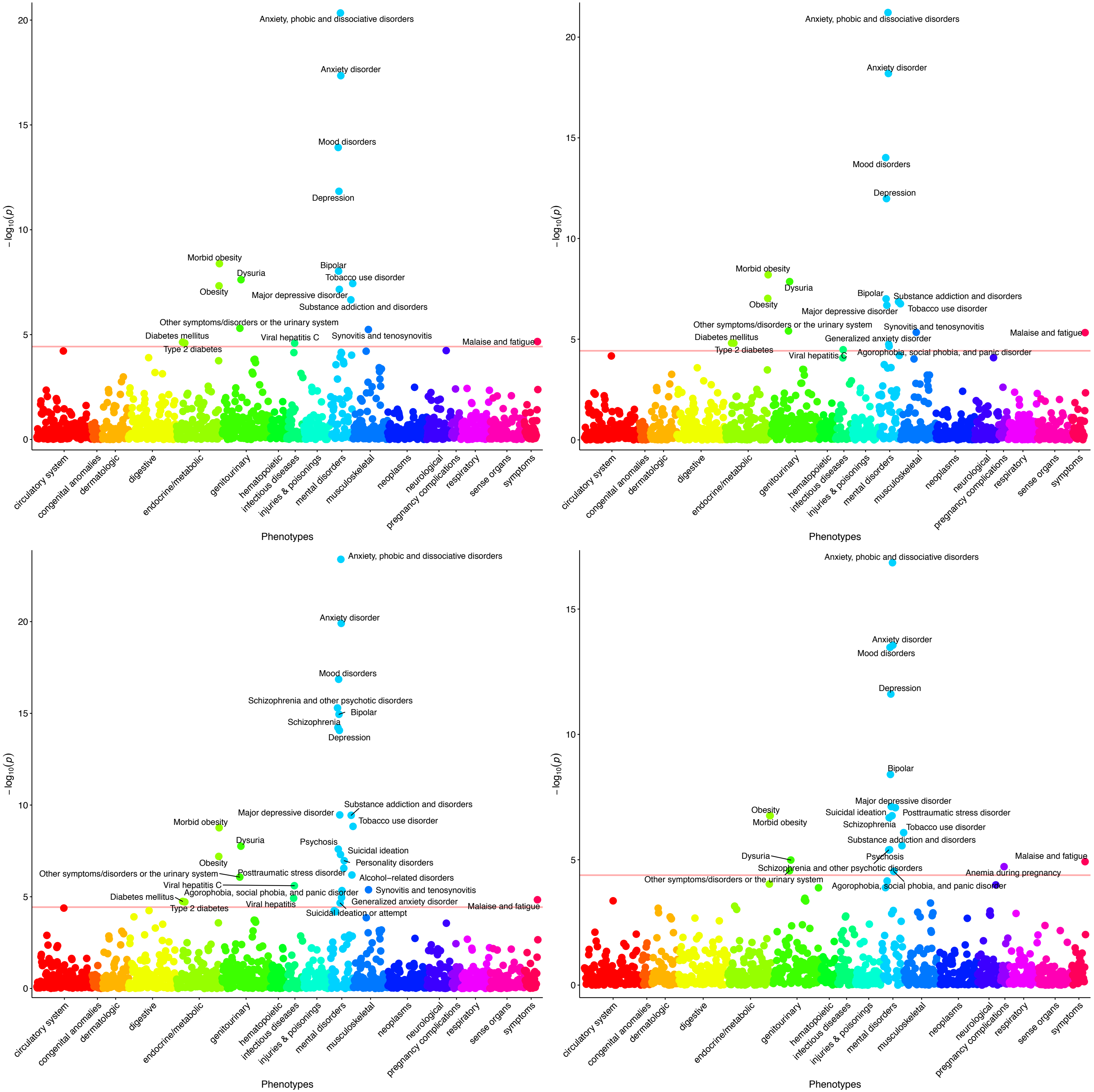
